# Directed evolution of *Anabaena variabilis* phenylalanine ammonia-lyase (PAL) identifies mutants with enhanced activities

**DOI:** 10.1101/2020.02.20.933945

**Authors:** Zachary JS Mays, Karishma Mohan, Vikas D Trivedi, Todd C Chappell, Nikhil U Nair

## Abstract

There is broad interest in engineering phenylalanine ammonia-lyase (PAL) for its biocatalytic applications in industry (fine-chemicals and natural product synthesis) and medicine (phenylketoruria/PKU and cancer treatment). While site-specific mutagenesis has been employed to improve PAL stability or substrate specificity, a more comprehensive mutational landscape has yet to be explored for this class of enzymes. Here, we report development of a directed evolution technique to engineer PAL enzymes. Central to this approach is a high-throughput enrichment that couples *E. coli* growth to PAL activity. Using the clinically-relevant PAL from *Anabaena variabilis*, which is used on the formulation of pegvaliase for PKU therapy, we identified mutations at residues previously unknown as relevant for function that increase turnover frequency almost twofold after only a single round of engineering. This work demonstrates the power our technique for ammonia-lyase enzyme engineering.

---

The ammonia lyase (AL; EC 4.3.1.x) class and aminomutase (AM; 5.4.3.x) class of enzymes have been the focus of decades of research and development for industrial and biomedical applications. Their prosthetic group, 4-methylideneimidazole-5-one (MIO), either catalyzes the transformation of an L-α-amino acid into the α,β-unsaturated carboxylic acid counterpart via the non-oxidative elimination of ammonia or into the spatially isometric β-amino acid via a 2→3 amine shift, respectively (Cooke et al., 2009). Accordingly, optically pure amino acids can be biosynthesized from the reverse reactions (D’Cunha et al., 1996; Shetty et al., 1986; Yamada et al., 1981), and the application of MIO-enzymes in both directions has yielded intermediates for pharmaceuticals (Lee et al., 2015; Walker et al., 2004), agrochemicals (An et al., 2007; Hoagland, 1996; Shin et al., 2012), polymers (McKenna and Nielsen, 2011; Verhoef et al., 2009, 2007), and flavonoids (Jiang et al., 2005; Lee et al., 2015; Park et al., 2012; Wu et al., 2013).

Phenylalanine ammonia lyase (PAL) has been of great interest as a treatment for the genetic disease phenylketonuria (PKU). Daily subcutaneous injection of a purified and PEGylated recombinant PAL from *Anabaena variabilis* (PEG-r*Av*PAL; Palynziq®, BioMarin Pharmaceutical Inc.) was approved by the US FDA in 2018 as an enzyme substitution therapy for PKU (Hydery and Coppenrath, 2019). Concurrently, an orally administered engineered probiotic *Escherichia coli* Nissle 1917 expressing recombinant PAL from *Photorhabdus luminescens* is currently under investigation by Synlogic Inc (Isabella et al., 2018). Other formulations of this enzyme are also being explored as therapeutics (Abell and Stith, 1973; Babich et al., 2013; Chang et al., 1995; Durrer et al., 2017; Rossi et al., 2014; Yang et al., 2019) as well as for the production of low phenylalanine (phe) protein dietary supplementation for PKU (Castañeda et al., 2015) and cancer (Kakkis et al., 2009; Shen et al., 1977) patients.

This significant interest has resulted in various efforts to improve enzyme properties. Structural and sequence homology between aromatic ALs and AMs has fuelled rational engineering efforts to alter or improve stability (Bell et al., 2017; Chesters et al., 2012; Wang et al., 2008; Zhang et al., 2017), substrate specificity (Bartsch et al., 2013; Louie et al., 2006; Lovelock and Turner, 2014; Xiang and Moore, 2005), and enantioselectivity (Turner, 2011; Wohlgemuth, 2010; Wu et al., 2009). However, application of combinatorial approaches that leverage evolutionary selection to search large sequence spaces for improved properties (Wrenbeck et al., 2017) has not been well-explored for this class of enzymes (Flachbart et al., 2019). Here, we developed a growth-based high-throughput enrichment scheme and screened a mutagenized PAL library to identify variants with improved kinetic properties. Core to this enrichment is the growth rescue of *E. coli* by PAL in minimal medium with phe, which cannot be used as the sole nitrogen source by K-12 strains (Reitzer, 1996). Consequently, *E. coli* can only grow if PAL actively deaminates phe to release ammonium, a highly preferred nitrogen-source (Figure 1a). We executed our directed evolution technique using the *A. variabilis* PAL (DM-r*Av*PAL/*Av*PAL*) (Kang et al., 2010; Sarkissian et al., 2008), because of its clinical significance, and identified mutants with improved catalytic properties. The mutations identified here have not been previously reported as important for PAL catalytic activity, demonstrating the advantage of our approach for scanning unexplored sequence space over previous efforts.

**Figure 1.**
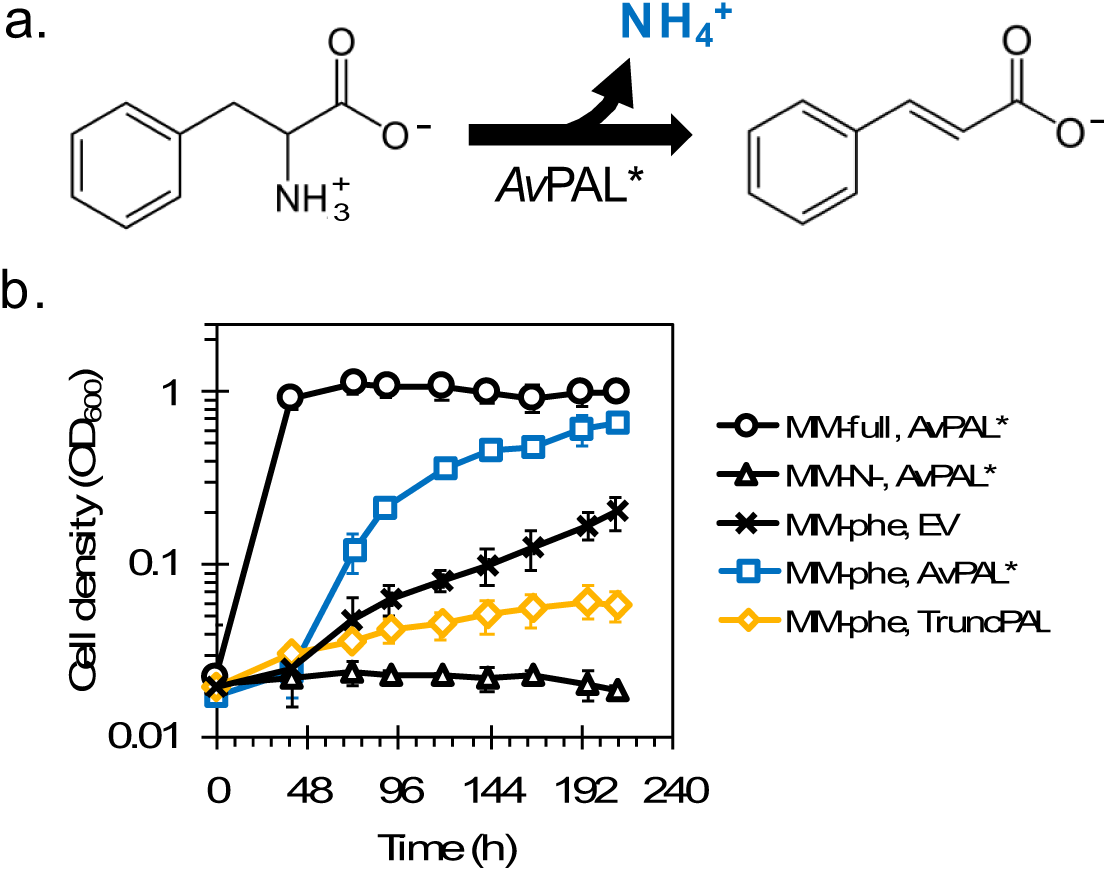
Initial study demonstrating growth-rescue of E. coli by PAL activity. (a.) Growth rescue relies on deamination of phenylalanine by PAL to form ammonium (NH_4_^+^), a preferred nitrogen source for E. coli. (b.) *E. coli* cells expressing active *Av*PAL* (□) in MM^phe,init^ grow faster than wild-type cells (×) or those expressing truncated inactive *Av*PAL* (◇). Cells grown in MM^full,init^ (○) and MM^N-,init^ (△) as controls.

Initial growth studies demonstrated that *Av*PAL* could rescue growth of *E. coli* in phe selective minimal media (MM^phe,init^) with a ∼70% biomass yield relative to complete minimal media (MM^full,init^), demonstrating coupling between growth to enzyme activity (Figure 1b). However, controls strains had unexpected, albeit slow, growth. Overexpression of noncatalytic proteins such as green fluorescent protein (sfGFP) or a truncated *Av*PAL* (TruncPAL) decreased background growth (Figure 1b) but still adversely affected the dynamic range to reliably select for highly active PAL over inactive mutants or other suppressors, if left unoptimized. Phenylalanine metabolism under austere conditions, *viz* nitrogen starvation, has not been well studied, and transaminases (AspC, IlvE, TyrB, HisC) may have unreported promiscuous activity on phe (Gelfand and Steinberg, 1977; Guzmán et al., 2015). Unfortunately, we observed no difference in the basal growth of *E. coli* in MM^phe,init^ after deleting each transaminase (Figure S1) suggesting no single gene was responsible for basal growth.

As an alternative approach to minimize accumulation of false positives, we optimized conditions to increase biomass yield and shorten the lag. We tested different media formulations (carbon source, pH, strain background, culture volume, phe concentration, and the presence of an additional nitrogen source) to achieve this goal (Figure 2a). We initially performed this optimization using MG1655, which grows poorly in minimal media because of inefficient pyrimidine utilization from a mutation in *rph* (Jensen, 1993). Switching to a strain with a corrected allele (MG1655^*rph+*^) shortened the lag phase by 18 h and culturing in glucose reduced the lag phase by another 24 h (Figure 2b).

**Figure 2.**
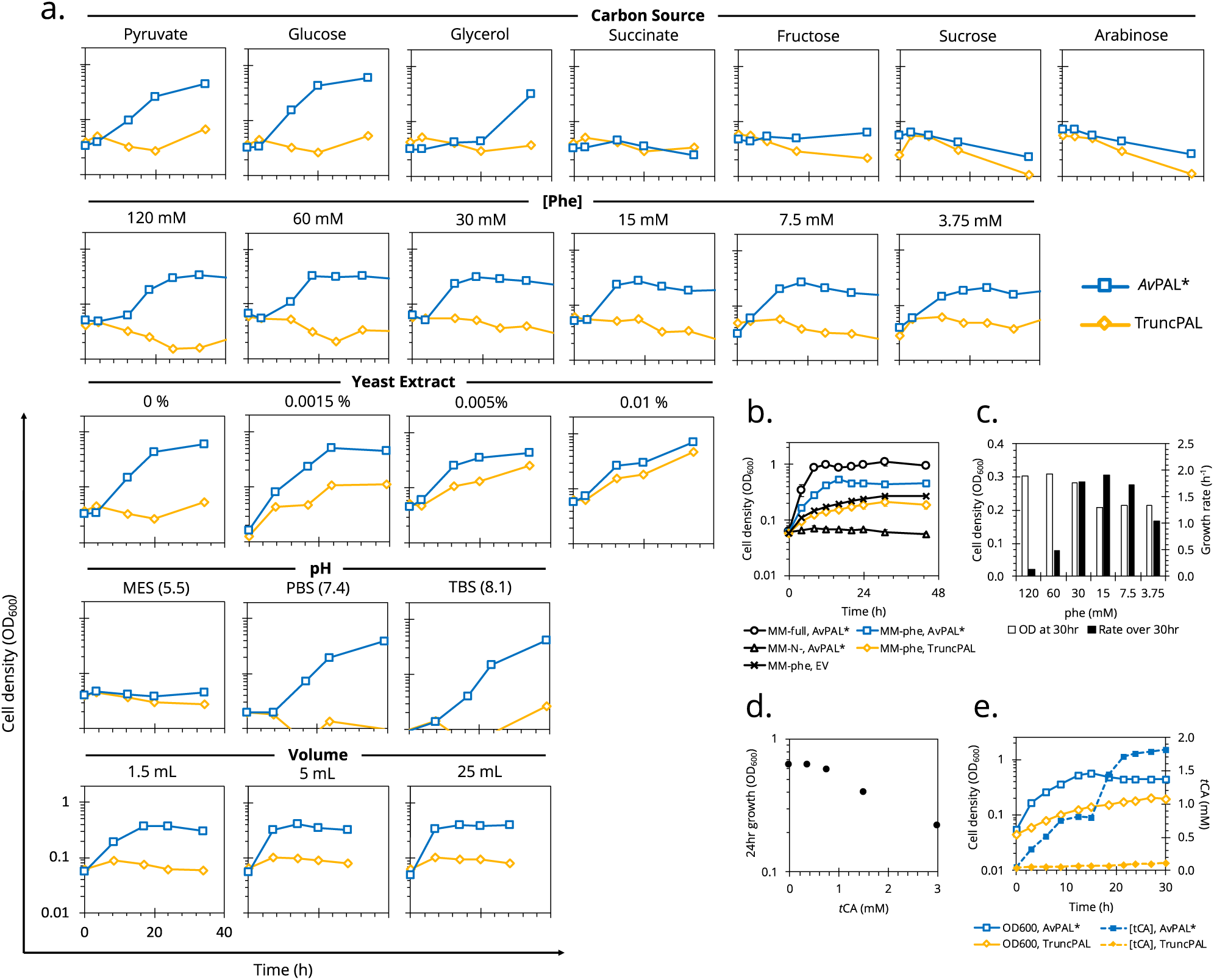
Optimizing conditions of growth-based PAL selection. (a.) *E. coli* MG1655(DE3)^*ΔendA,ΔrecA*^ cells expressing *Av*PAL* (blue) or truncated *Av*PAL* (orange) were tested for growth in MM^phe^ under different conditions. (b.) The final optimized conditions allowed for growth to be recovered in 12 h compared to 3 d previously. (c.) While optimizing the growth media, we observed that at phe concentrations > 30 mM, final biomass density decreased and lag time increased, suggesting toxicity due to rapid accumulation of *t*CA. At phe concentrations < 30 mM, final biomass densities dropped, and at concentrations ≤ 7.5 mM, growth rate was also slowed, suggesting insufficient nitrogen to sustain growth. (d.) Media supplemented with *t*CA inhibited the growth of *E coli* at concentrations ≥ 1 mM. (e.) *Av*PAL* expressing cells produce and secrete *t*CA to ∼ 1 mM tCA before arresting growth. Subsequent bolus increase in tCA during death phase is likely due to cell lysis. All curves representative of duplicates with less than 10% error.

We found that not only was phe concentration important for optimal growth (Figure 2a,c), but that *trans*-cinnamic acid (*t*CA) was toxic to cells. Cells grown in MM^full,opt^ showed impaired growth when supplemented with 1.5 mM tCA (Figure 2c). When grown in MM^phe,opt^ containing <30 mM phe, growth rate and biomass yield were reduced by low nitrogen availability. However, at phe >30 mM, *t*CA accumulated too quickly causing toxicity, and the cells not only experience a long lag but also quickly arrest growth (Figure 2c-e). The final optimized conditions in Figure 2e show that despite growth levelling out at OD_600_ 0.3, the lag was virtually eliminated. Thus, we determined that subculturing the cells into fresh medium at OD_600_ 0.2 would minimize *t*CA toxicity and basal growth—maximizing the difference between inactive and active PAL expressing cells. To validate enrichment occurs under these conditions, we created a mock library by transforming a 1:10 or 1:1000 mixture of *Av*PAL*-to-sfGFP-expressing. We measured cell fluorescence by flow-cytometry and observed decreasing fluorescence and increasing PAL activity at successive rounds of subculture in MM^phe,opt^ (Figure S2).

After finalizing the conditions to enrich active PAL, we created a mutant library of 10^5^ variants with an average error rate of 2.8 aa/protein. The entire library was grown in MM^phe,opt^ over three rounds, subculturing each time at OD_600_ of 0.2 (Figure 3a). We subsequently plated the cells on non-selective LB medium, picked five random colonies and screened their lysates for PAL activity. Two of the five, *viz* M222L and L4P/G218S, showed nearly twofold higher activity with the other three showing similar activity to parental *Av*PAL* (Figure 3b). This result suggests successful enrichment of higher activity mutants over lower/inactive mutants. *E. coli* expressing M222L and L4P/G218S mutants showed faster growth compared to *Av*PAL* in MM^phe,opt^, with all attaining the same OD_600_ at stationary phase (Figure 3c). The greater differences in growth profiles at early growth stages between mutants (M222L and L4P/G218S) and parental *Av*PAL* is consistent with the enrichment strategy of subculturing at low OD_600_. Furthermore, residues 218 and 222 are directly adjacent to the active site of *Av*PAL* and in close vicinity of the MIO-adduct (Wang et al., 2008). Comparing the crystal structure of *Av*PAL* to these mutants shows potential changes in hydrogen bonding within the active site (Figure S3).

**Figure 3.**
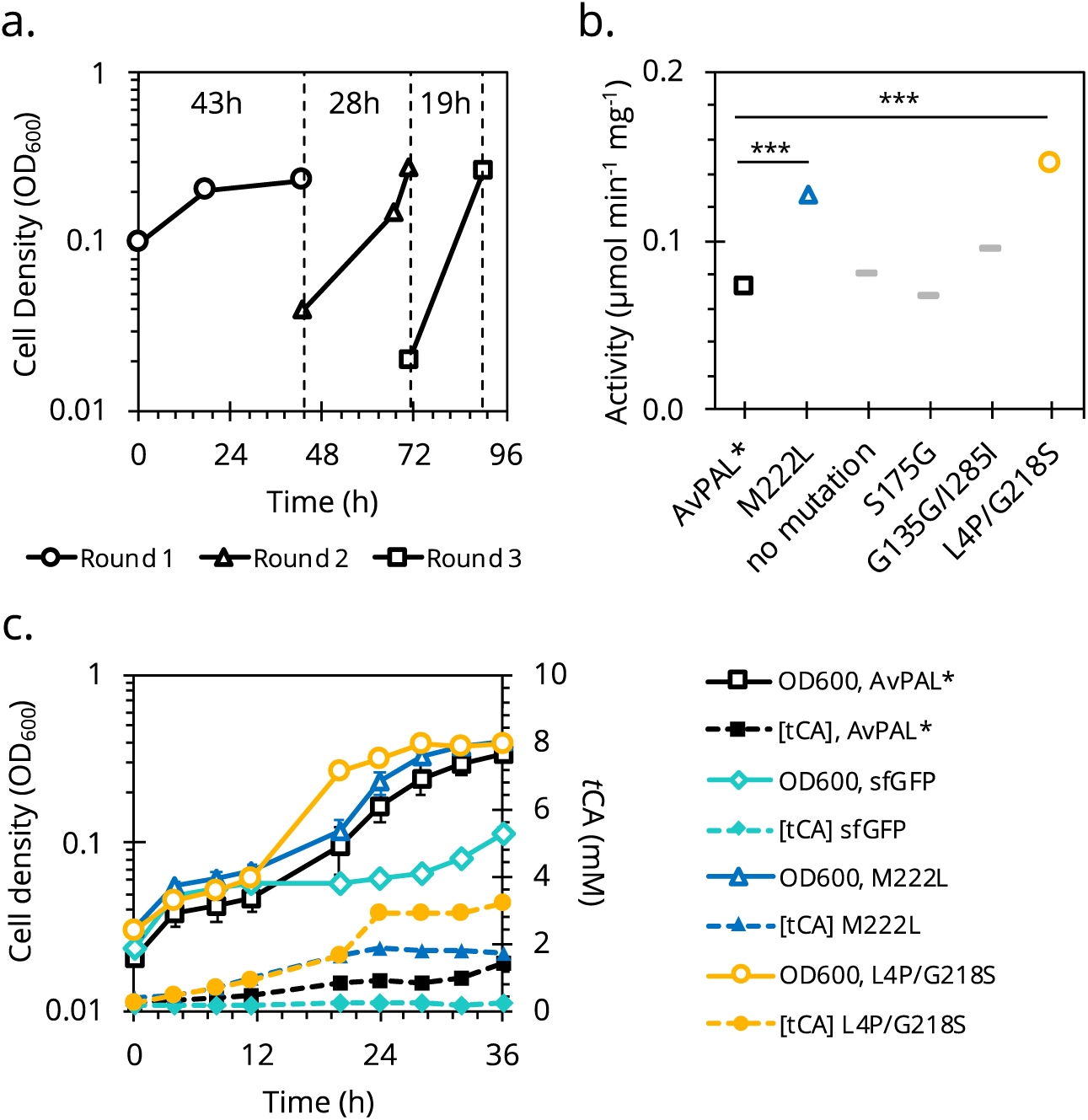
Identification of *Av*PAL* mutants by growth-coupled enrichment. (a.) The growth profiles of *E. coli* MG1655^*rph+*^ cells expressing the *Av*PAL* mutant library grown in MM^phe,opt^ over three rounds. (b.) PAL activity in lysate of 5 randomly picked colonies normalized to total protein. (c.) The growth profiles (solid) and *t*CA production (dotted) in MM^phe,opt^ of mutants showing higher than parental activity.

Previous studies with *Av*PAL* have demonstrated that kinetic parameters, pH optimum, thermal and proteolytic stabilities are relevant to therapeutic efficacy for PKU enzyme-replacement therapy. The k_cat_ of both the mutants was 70 - 80% higher than parental *Av*PAL* (Figure 4a) whereas the K_M_ of M222L was similar to that of the parent and that of L4P/G218S was ∼2.5× higher. Overall, the M222L mutant showed improved catalytic efficiency compared to *Av*PAL*, while L4P/G218S mutant showed a trade-off between turnover frequency and substrate “affinity”. *Av*PAL* is reported to have a pH optimum in the range of 7.5 - 8.5 (Wang et al., 2008) and we observed similar results for both the mutants (Figure 4b), albeit with a slightly narrower optimal range. Temperature stability was assessed by subjecting the mutants to different temperatures for 1 h before measuring enzyme activity at optimal conditions (37 °C, pH 7.4). The enzymes remained stable from 37 °C to 55 °C and began a modest decrease in relative activity at 65 °C before denaturing at 80 °C (Figure 4c). The proteolytic stability was evaluated by incubating purified enzymes to trypsin. M222L was as trypsin-resistant as *Av*PAL* but L4P/G218S showed rapid loss of activity within five minutes (Figure 4d).

**Figure 4.**
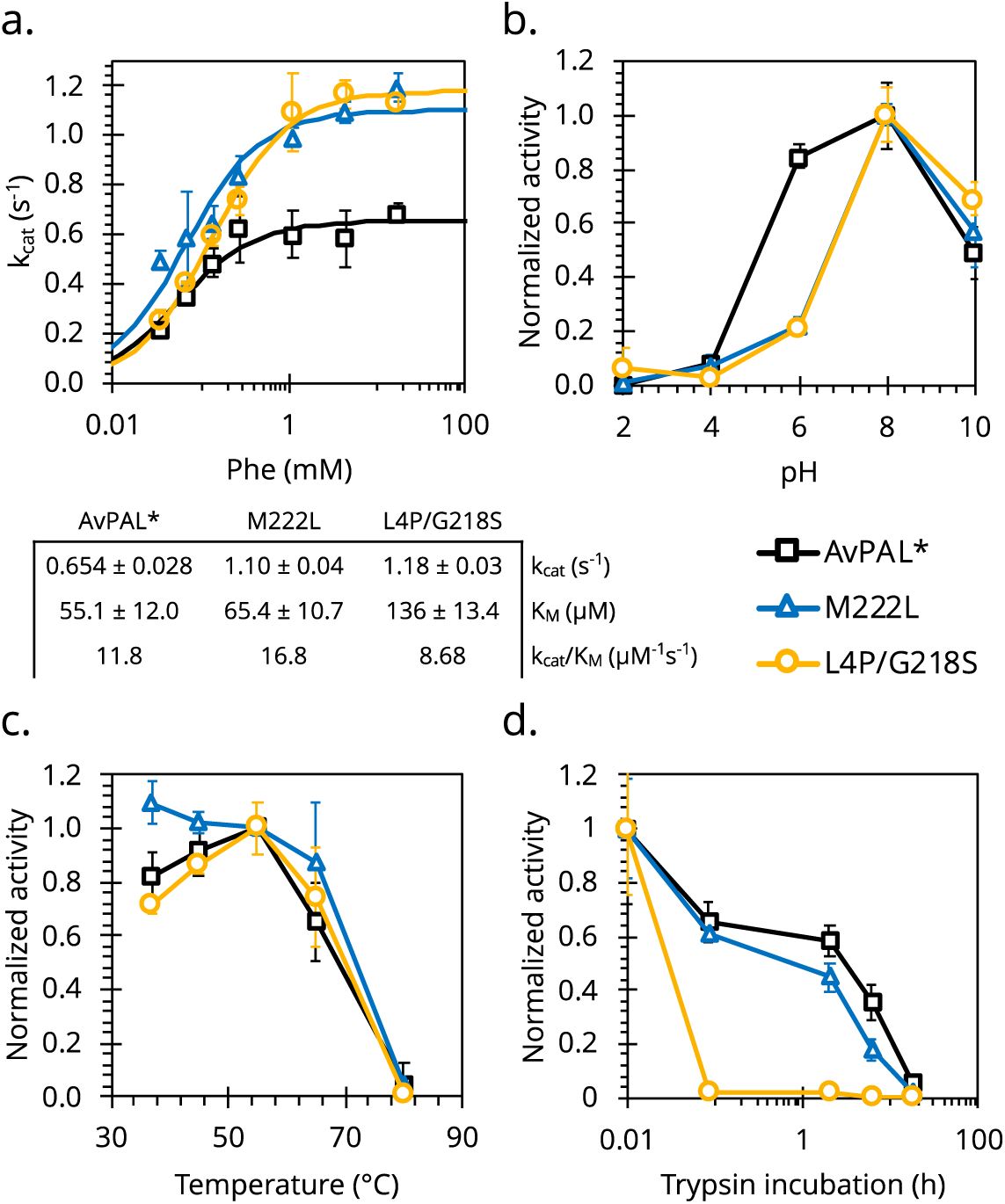
Biochemical characterization of PAL mutants. Two mutants showing higher than wildtype activity were characterized to establish (a.) kinetic parameters, (b.) pH optimum, (c.) temperature optimum, (d.) and resistance to protease degradation.

Our results show that the catalytic properties of this class of enzymes, which are important for both industrial and biomedical applications, can be engineered using directed evolution. Further, the large sequence space we rapidly searched to identify mutations at residues previously unrecognized as functionally important, serves as evidence of this technique’s strength. Since deamination activity serves as the foundation of technique, we offer this method as demonstration that may be applicable to other ammonia lyases as well. The improvements in turnover rates observed here are unprecedented in the literature, either through rational or combinatorial methods, and has tremendous translation potential, especially for PKU.

## Material & Methods

### Microbial strains, plasmids, and growth conditions

All *Escherichia coli* strains were cultured in lysogeny broth (LB) (VWR International, Randor, PA) at 37 °C with rotary shaking at 250 rpm. All media was solidified using 1.5 % (w/v) agar (Teknova Inc, Hollister, CA). Minimal media (MM) conditions are described in the “Optimization of growth-coupled enrichment” section below. *E. coli* DH5α was used as a host for the construction of the expression vectors and cultured as above only supplemented with chloramphenicol (25 µg/mL) (RPI Corp). Initial expression in MM was performed in *E. coli* MG1655(DE3)^*ΔendA,ΔrecA*^ and later moved to *E. coli* MG1655^*rph+*^ for final experiments.

All cloning was performed in *E. coli* DH5α with reagents from New England Biolabs, Inc (Ipswich, MA). Preliminary expression experiments were conducted using the inducible pACYC-Duet1_*Av*PAL*, constructed by using the surrounding sites for restriction endonucleases NcoI and XhoI. For subsequent experiments requiring constitutive expression, the plasmid pBAV1k was implemented, with the gene of interest was replaced with *Av*PAL* using Gibson assembly.

### Enzyme activity assays

The activity of all *Av*PAL constructs was measured by production of *t*CA over time. Cultures were sonicated on ice using a Sonifier SFX 150 (Branson Ultrasonics, Danbury, CT) (2 s on; 10 s off; 4 min; 55%), and debris was separated from the lysate by centrifuging at 10,000 × g for 10 min. Ten microliters of lysate were then mixed with 190 μL of pre-warmed 50 mM phe (Tokyo Chemical Industry, Portland, OR) in phosphate buffered saline (PBS) (137 mM NaCl, 2.7 mM KCl, 10 mM Na_2_HPO_4_, 2 mM KH_2_PO_4_, pH 7.4) in a 96-well F-bottom UVStar (Greiner Bio-One, Kremsmünster, Austria) microtiter plate. Absorbance at 290nm was measured every 30 s at 37°C using a SpectraMax M3 (Molecular Devices) plate reader.

Each construct included a N-term His-tag used for immobilized metal affinity chromatography (IMAC) purification. Briefly, overnight cell cultures were sonicated in 3 mL Equilibration buffer (300 mM NaCl, 50 mM NaH_2_PO_4_, 10 mM imidazole, 15% (w/v) glycerol, pH 8.0). The lysate was loaded onto a prepared column with 2 mL TALON Metal Affinity Resin (Clontech Laboratories, Inc., Mountain View, CA). After being washed twice with 5 column volumes (CV) of Equilibration buffer, pure protein was then eluted off the column with 2.5 mL of Elution buffer (300 mM NaCl, 50 mM NaH_2_PO_4_, 500 mM imidazole, 15% (w/v) glycerol, pH 8.0), collecting 0.5 CV fractions until dry. Elution fractions showing clean protein bands on an SDS-PAGE were then dialyzed and concentrated in Storage buffer (20% (v/v) glycerol in PBS, pH 7.4) using a 10K MWCO Microsep Advance Centrifugal Device (Pall Corporation, Port Washington, NY) as directed. Purified protein extracts were aliquoted and stored at -20 °C, replacing lysate in subsequent characterization and activity assays. Protein concentration was measured by Bradford method using bovine serum albumin (BSA) as the standard.

### *Av*PAL library creation

Random mutagenesis libraries were created using two rounds of error prone PCR, with the amplicon of the first reaction serving as the template DNA for the second. Each reaction contained 1X Standard *Taq* reaction buffer (New England Biolabs, Inc.), 5 mM MgCl_2_, 0.15 mM MnCl_2_, 0.2 mM dATP, 0.2 mM dGTP, 1 mM dCTP, 1 mM dTTP, 0.4 µM each primer, 0.4 ng/μl template DNA, and 0.05 U/ml *Taq* DNA polymerase. The reactions were amplified using the following PCR cycle conditions: 95 °C denaturation, 1 min; 16 cycles of 95 °C denaturation, 30 s; 46 °C annealing, 45 s; and 68 °C extension, 2 min, followed by 68 °C extension for 5 min. The target vector, pBAV1k was amplified separately using *Phusion* PCR, and the two were combined using Gibson assembly. The reaction was purified with a E.Z.N.A. Cycle Pure Kit (Omega) before being transformed by electroporation into *E. coli* MG1655^*rph+*^.

### Optimization of growth-coupled enrichment

Growth was measured by seeding cultures at OD_600_ 0.05 and monitoring cell density over time. Initial experiments used a base nitrogen-deficient minimal media (MM^N-^) (33.7 mM Na_2_HPO_4_, 22 mM KH_2_PO_4_, 8.55 mM NaCl, 1 mM MgSO_4_, 0.1 mM CaCl_2_, 10 μM FeSO_4_, 0.4% (v/v) glycerol, 10 μg/mL thiamine, 20 μM IPTG, 12.5 μg/mL chloramphenicol, pH 7.4) that was supplemented with 9.35 mM phe (MM^phe,init^) or 9.35 mM NH_4_Cl (MM^full,init^). Variable conditions were changed across the parameters outlined in Figure 2, as well as moving to a more favorable strain for growth in minimal media. This resulted in a final MM^N-,opt^ (137 mM NaCl, 2.7 mM KCl, 10 mM Na_2_HPO_4_, 2 mM KH_2_PO_4_, 2 mM MgSO_4_, 1x Trace Metals (Teknova, Inc.), 0.2% (v/v) glucose, 10 μg/mL thiamine, 12.5 μg/mL chloramphenicol, pH 7.4) supplemented with 30mM phe (MM^phe,opt^) or 9.35 mM NH_4_Cl (MM^full,opt^). To enrich the active population of the *Av*PAL* library, cells were subcultured into fresh MM^phe,opt^ once they reached OD_600_ 0.2. Remaining cells in each round were miniprepped as a pooled plasmid library for further analysis.

### Flow cytometry

Plasmids, both with a pBAV1k backbone, expressing sfGFP or *Av*PAL* were mixed in a 1000:1 or 10:1 ratio as a mock mutant library and transformed by electroporation into *E. coli* MG1655^*rph+*^. Cells were recovered for 1 h before being washed and seeded in 5 mL of selective media as prepared above. Cell density was measured over time until reaching OD_600_ 0.2, when the cells were subcultured to OD_600_ 0.05 for the next round of enrichment. Cells were also plated at each subculture for PCR amplification to confirm the presence of either sfGFP or *Av*PAL*. Cells at each point of subculture were also diluted to OD_600_ 0.05 for flow cytometry analysis. A minimum of 10,000 events were collected using a blue laser on an Attune NxT flow cytometer (Life Technologies, Carlsbad, CA). Fluorescence of sfGFP was detected on the BL1-H channel with 488nm excitation, and loss of fluorescence was revealed as a measure of active *Av*PAL* enrichment.

### Enzyme kinetics

*Av*PAL* and selected mutants were purified as described above. The activity of 0.1 µg of protein was measured by the production of *t*CA over 10 min by recording the absorbance of t ahe reaction mix at 290nm. Phe was added at varying concentrations from 35µM to 17.5mM in PBS, pH 7.4 (PBS) at 37°C to begin the reaction. A Michaelis-Menten curve was fit in GraphPad Prism software using the initial rate at each phe concentration.

### pH profile

The optimal pH of *Av*PAL* and selected mutants was determined by performing the enzyme activity described above. A 35 mM phe solution was buffered across a pH range (2 to 10) using phosphate - citrate buffer, prepared by varying concentrations of Na_2_HPO_4_ and citric acid. Total 0.2 µg protein was used to carry out the activity reaction in 200µl at 37°C.

### Temperature stability

The effect of temperature on the stability of *Av*PAL* and selected mutants was determined by incubating the protein in PBS, pH 7.4 at temperatures ranging from 37°C to 80 °C for 1 hour followed by measuring the enzyme activity. Each enzyme reaction was carried out using 1 µg of PAL protein and 35mM phe as substrate in a total reaction volume of 200µl at 37 °C.

### Proteolytic stability

The proteolytic stability was evaluated by subjecting *Av*PAL* and selected mutants to a catalytic amount of trypsin as previously described^36^. Briefly, 100 µg/ml *Av*PAL enzyme was subjected to trypsin (40 µg/ml) (MilliporeSigma, Burlington, MA) in PBS at 37 °C. Enzyme activity of 1 µg of protein was then measured as described above.

## Acknowledgments

We would like to thank Prof. Nicholas Turner (University of Manchester, UK) for sharing the *Av*PAL plasmid. This work was supported by NIH grants DP2HD91798 and R03HD090444.

