## Supplemental Information for "Directed evolution of *Anabaena variabilis* phenylalanine ammonia-lyase (PAL) identifies mutants with enhanced activities"

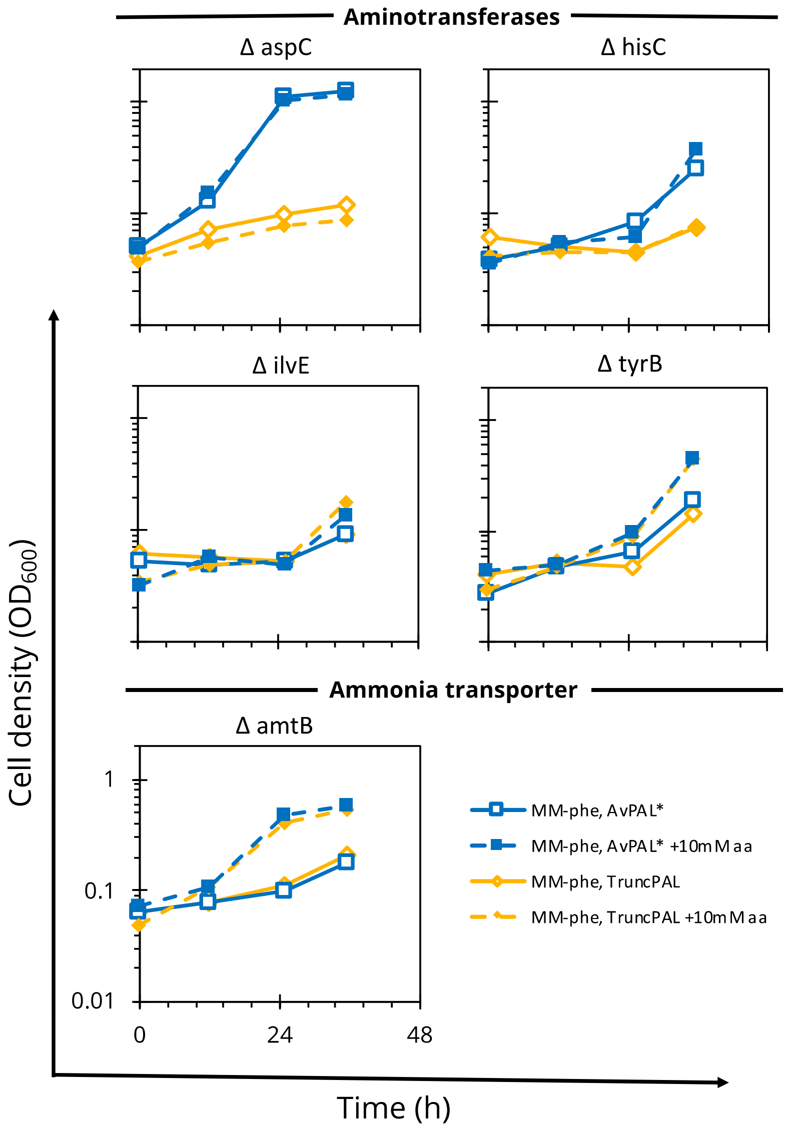


**Figure S1.** Growth of *E. coli* after gene deletions intended to lower basal growth on $MM^{phe,init}$. (a.-d.) Select aminotransferases with reported promiscuous activity on phenylalanine were deleted in an attempt to reduce the level of basal growth seen by wild-type *E. coli* on $MM^{phe,init}$. Each deletion strain showed no changes in growth whether or not expressing AvPAL*. (e.) The ammonia transporter AmtB was also deleted in an attempt to minimize cross-feeding of nitrogen between cells but had no benefit.

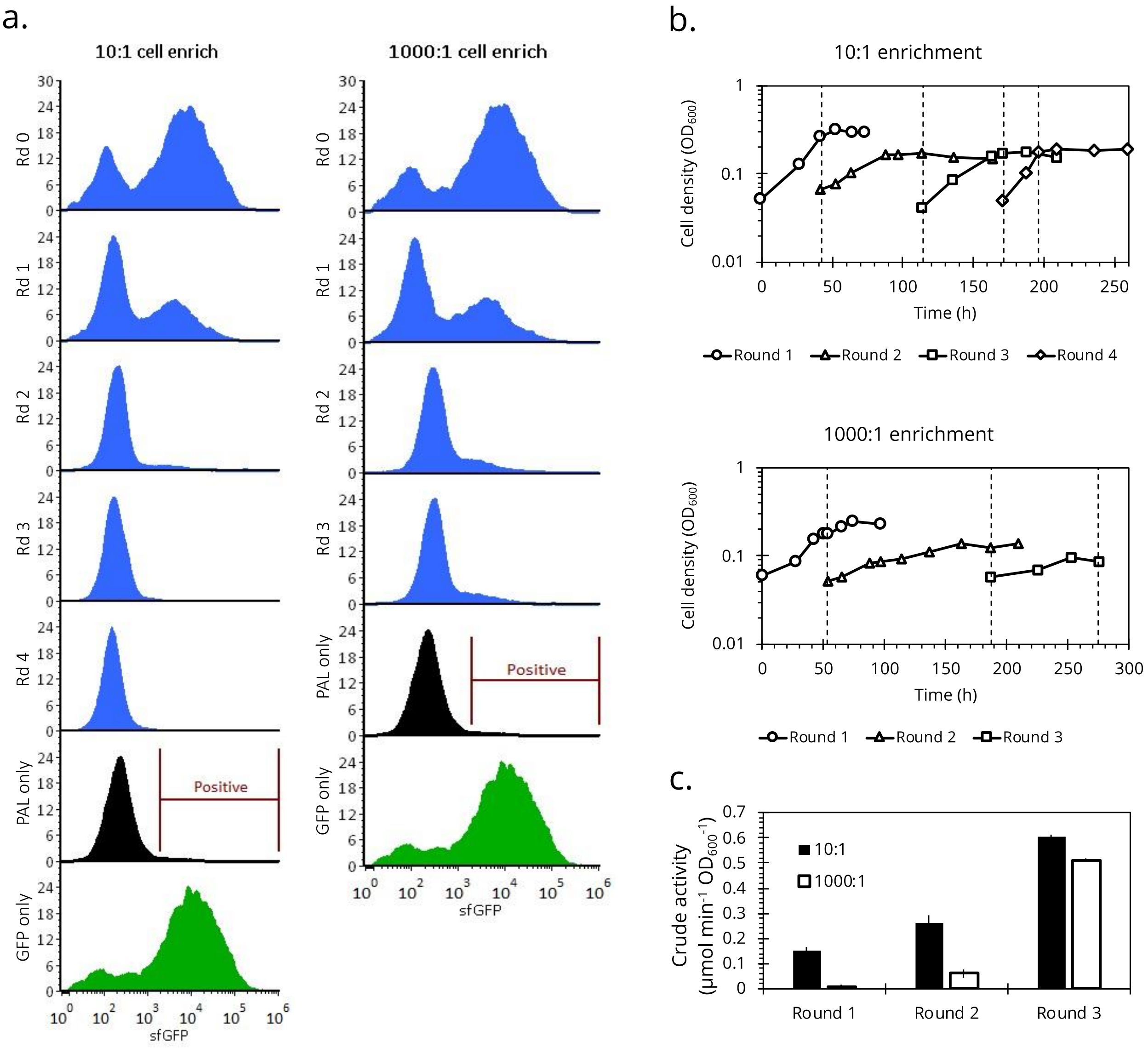


**Figure S2.** Validating enrichment with a mock library. After transforming a plasmid mix of *Av*PAL* and sfGFP in 1:10 or 1:1000 ratio, we were able to observe (a.) the loss of fluorescence, and (b.) the enrichment of cells expressing *Av*PAL* over sfGFP, over rounds of subculturing in $MM^{phe}$ selective media. This was confirmed by (c.) an observed increase in *Av*PAL* activity on a per cell basis.

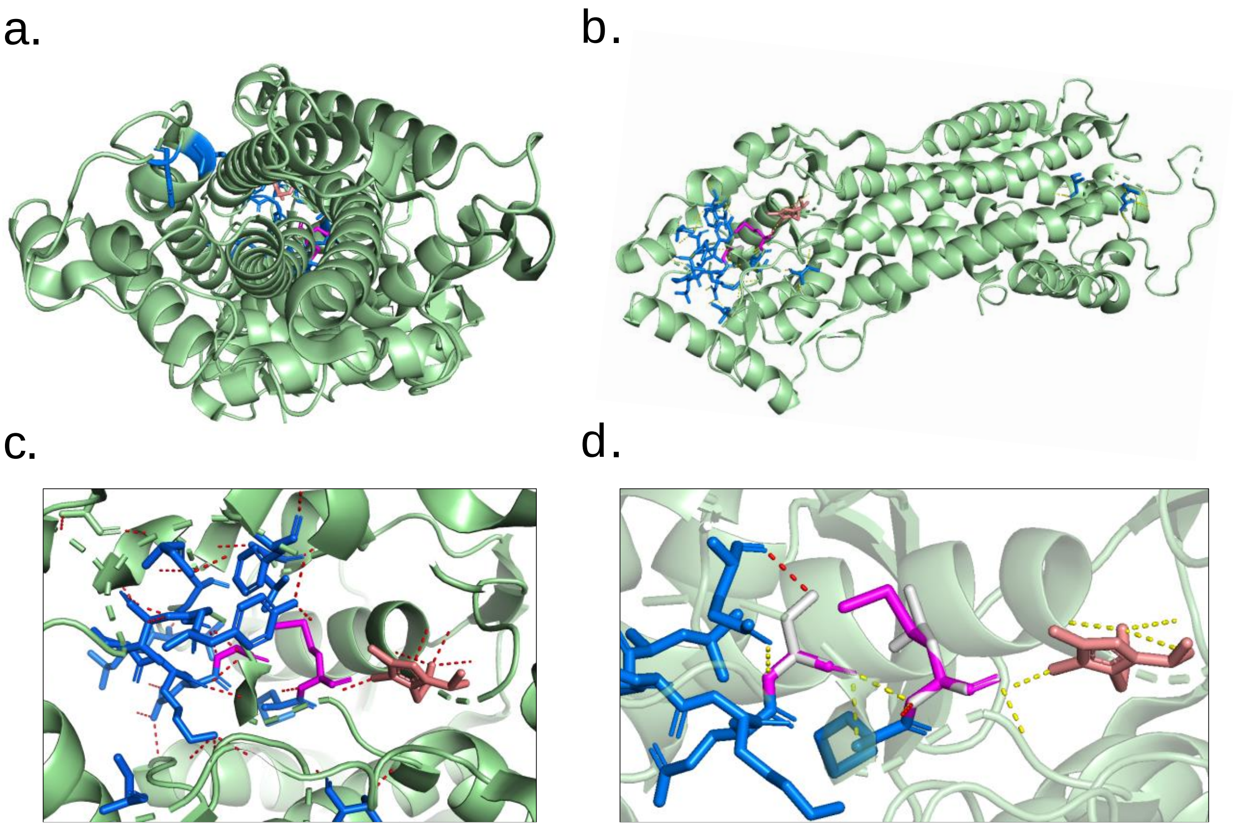


**Figure S3.** Crystal structure analysis of *Av*PAL* monomer with active site residues (blue), MIO-adduct (orange), and residues 218 and 222 (pink) highlighted. (a.) A top view looking down into the active site. (b.) Side-view of the monomer. (c.) Close up of the wildtype *Av*PAL* active site with predicted intra-residue hydrogen bonding. (d). Comparison of the wildtype and mutant active sites with residues 218 (left) and 222 (right) highlighted. Mutant residues G218S and M222L (gray) have altered intra-residue hydrogen bonding (red, dotted) compared to wildtype (yellow, dotted).
